# Rapid geometric feature signaling in the spiking activity of a complete population of tactile nerve fibers

**DOI:** 10.1101/461095

**Authors:** Benoit P. Delhaye, Xinyue Xia, Sliman J. Bensmaia

## Abstract

Tactile feature extraction is essential to guide the dexterous manipulation of objects. The longstanding theory is that geometric features at each location of contact between hand and object are extracted from the spatial layout of the response of populations of tactile nerve fibers. However, recent evidence suggests that some features (edge orientation, e.g.) are extracted very rapidly (<200ms), casting doubt that this information relies on a spatial code, which ostensibly requires integrating responses over time. An alternative hypothesis is that orientation is conveyed in precise temporal spiking patterns. Here, we simulate, using a recently developed and validated model, the responses of tactile fibers from the entire human fingertip (∼800 afferents) to edges indented into the skin. We show that edge orientation can be quickly (<50 ms) and accurately (<3°) decoded from the spatial pattern of activation across the afferent population, starting with the very first spike. Next, we implement a biomimetic decoder of edge orientation, consisting of a bank of oriented Gabor filters, designed to mimic the documented responses of cortical neurons. We find that the biomimetic approach leads to orientation decoding performance that approaches the limit set by optimal decoders and is actually more robust to changes in other stimulus features. Finally, we show that orientation signals, measured from single units in non-human primate cortex (2 macaque monkeys, 1 female), follow a time course consistent with that of their counterparts in the nerve. We conclude that a spatial code is fast and accurate enough to support object manipulation.

## Introduction

Our sense of touch plays an important role in guiding even simple interactions with objects – such as picking up a key off a flat surface – and is essential for fine dexterous manipulation – such as threading a needle or clasping a necklace. This dexterity relies on precise tactile signals conveyed by mechanosensitive nerve fibers that innervate the fingertips (Johansson and Flanagan, 2009). An important object feature to be signaled by these populations of nerve fibers is local shape. For example, grasping a mobile phone requires sensing the location and orientation of its edges at each point of contact between hand and phone. Object shape was long thought to be extracted from a spatial pattern of activation in tactile nerve fibers (Phillips and Johnson, 1981; LaMotte and Srinivasan, 1987a, 1987b; Srinivasan and LaMotte, 1987; Johnson and Hsiao, 1992; LaMotte et al., 1996; Dodson et al., 1998), culminating in orientation tuned responses of neurons in somatosensory cortex (Pubols and Leroy, 1977; DiCarlo and Johnson, 2000; Hsiao et al., 2002; Bensmaia et al., 2008a; Delhaye et al., 2018). Orientation tuning in somatosensory cortical neurons emerges from the spatial layout of their receptive fields (RFs), which consist of excitatory fields flanked by inhibitory ones (DiCarlo et al., 1998; DiCarlo and Johnson, 1999, 2000, 2002), drawing strong analogies with early visual shape processing (Thorpe et al., 2001; Yau et al., 2009; Pack and Bensmaia, 2015).

That this process of feature extraction plays a role in object manipulation has been called into question, however. Indeed, for tactile signals to be effectively integrated into the planning and execution of the hand movements involved in object interactions, these signals need not only be precise but also rapid. The rapidity with which orientation information becomes available was demonstrated in a study that showed that human participants could align a tactile edge to a visual one with high accuracy (∼5 degrees), and that this orienting response occurred within 200 ms (Pruszynski et al., 2018). Whether edge orientation could be extracted so rapidly from a spatial pattern of activation, which presumably requires integrating signals over time, seemed dubious. Rather, information about shape was proposed to be encoded in the relative latencies of first spikes across populations of tactile fibers, or in the precise temporal pattern of spiking in individual fibers. Indeed, given the exquisite temporal precision of afferents spiking, such signals are in principle highly informative about shape (Johansson and Birznieks, 2004; Johansson and Flanagan, 2009; Pruszynski and Johansson, 2014).

Whether spike timing-based shape information can be reliably decoded has not been established, however, though some theoretical work in vision has revealed its potential (Van Rullen et al., 2005). In fact, temporal spiking patterns tend to be highly sensitive to the precise geometry and location of a stimulus, and orientation decoders built from temporal patterns in single afferents generalize poorly (Suresh et al., 2016). Furthermore, the time course of the spatial orientation signals has not been investigated – the aforementioned studies investigated only time-averaged responses –, leaving open the possibility that these signals may be sufficiently rapid to support object interactions.

In this study, we fill this gap by simulating – using a recently developed model that reconstructs afferent response with unprecedented spatial and temporal precision – the responses to indented edges of all the tactile fibers that innervate the glabrous skin (using TouchSim)(Saal et al., 2017). We then assess the time course over which information about orientation becomes available across this population response. We show that orientation can be accurately and rapidly decoded from the spatial pattern of activation, even with a mechanism inspired by the known RF properties of cortical neurons. We further demonstrate that taking into consideration precise spike timing does not substantially improve orientation acuity and does so at the expense of robustness to noise. Finally, we show that orientation signals in cortex follow a time course consistent with that of their counterparts in the nerve after accounting for conduction delays.

## Methods

### Population responses in the somatosensory nerves

#### Simulations

**Stimuli**. Simulated edges were indented into the skin at the center of the fingerpad at thirty equally spaced orientations ranging from 0 to 45 degrees (**Figure 2 A-B**). The rectangular edges were 1.6-mm wide, with aspect ratios ranging from 2 to 6 (3.2, 4.8, 6.4, 8, or 9.6-mm in length) and indentation depths ranging from 0.8 to 2.4 mm (in 0.4-mm increments), yielding a total of 3750 stimuli (30 orientation x 5 aspect ratios x 5 depths, each repeated 5 times). The indentation trajectory followed a trapezoidal profile and lasted 400 ms in total, including 50-ms constant speed on and off ramps and a 300-ms hold phase. To render the decoding problem more biologically relevant, we supplemented the intrinsic noise in the simulated afferents (Saal et al., 2017) with trial-to-trial variability in the form of a jitter in the position of the stimulus to mimic small variations in the stimulus presentations and small movements of the skin. The jitter consisted of isotropic, Gaussian noise, with mean zero, at one of four standard deviations (0, 0.25, 0.5 and 1 mm) to assess decoding robustness.

**Afferent responses**. We simulated the spiking responses of every tactile fiber that innervates the fingertip using a recently developed model, dubbed TouchSim (Saal et al., 2017). In brief, the model consists of a skin mechanics model – which characterizes how the skin deforms when a spatio-temporal stimulus is applied to its surfaced – and an integrate-and-fire model, which generates the spiking response of tactile nerve fibers in response to the skin deformation. Models corresponding to each afferent type tile the skin at their respective densities, yielding approximately 300 slowly adapting type I and 600 rapidly adapting fibers (cf. (Johansson and Vallbo, 1979))(**Figure 2C**). Pacinian (PC) afferents were not included in the simulations and analyses because their low density and poor spatial modulation makes them ill-suited to convey information about edge orientation (Saal et al., 2017). Slowly adapting type 2 (SA2) were not included because they are not included in TouchSim; while these fibers also likely convey information about edge orientation, these signals are probably less acute than are their SA1 and RA counterparts, given the respective densities and resulting spatial resolutions. TouchSim reproduces measured spiking responses with millisecond precision and most of the known response properties of tactile fibers (as detailed in (Saal et al., 2017)). The model is well suited to estimate responses to indented stimuli, even though the spiking models were fit mechanical noise comprising frequencies ranging from 1 to 1000 Hz. In the present simulations, conduction velocities were ignored so spikes were simulated as if they were measured as they are generated.

#### Validation of the simulation

TouchSim has been extensively tested and validated, with stimuli that include a variety of indented patterns (Saal et al., 2017). Nevertheless, we further validated the model by comparing simulated responses to indented edges to their measured counterparts (from previously published data, see (Bensmaia et al., 2008a)).

**Stimuli**. Stimuli were delivered with a dense array stimulator, consisting of 400 independently controlled pins arranged in a 10 mm x 10 mm grid (with a spacing of 0.5 mm) (Killebrew et al., 2007). On each trial, am edge was indented into the skin and held for 30 ms at one of eight orientations, ranging from 0 to 157.5° in steps of 22.5°, with 0° defined as the orientation perpendicular to the long axis of the finger. The duration of the on ramp was 20 ms, the depth of indentation (edge amplitude) was 500 µm, and the edge width was 1 mm. The stimulation parameters (indentation velocity, depth and edge length) were thus similar to the ones used in the peripheral nerve simulations. The pivot of the bar, which was also its center, was either located at the point of maximum sensitivity (or hotspot) of the nerve fiber or shifted by 1 to 5 mm (steps of 1 mm) along the axis perpendicular to the edge at each orientation. The inter-stimulus interval was 100 ms. Edges were each presented 10 times in pseudorandom order for a total of 880 trials (8 orientations x 11 offsets x 10 presentations).

**Neurophysiology**. Single unit recordings were obtained from the median and ulnar nerves of four anaesthetized macaque monkeys using standard methods (Mountcastle et al., 1967; Talbot et al., 1968) as previously described in details (Bensmaia et al., 2008a). Recording were made from afferents located on the distal fingerpads of digits 2-5. The afferent types were identified based on the standard criteria: afferent with small and well defined RFs were classified as SA1 (slowly adapting type 1) afferents if they evoked sustained firing during steady indentation or as RA (rapidly adapting) afferents if they only responded during movement. The point of maximal sensitivity was located using a handheld probe.

**Comparison of simulated and measured responses**. Using Touchsim, we simulated the responses of individual afferents (4 SA1 and 9 RA) to stimuli identical to those used in the neurophysiological experiments described above. We then compared the typical profiles of simulated and measured afferent responses. Given that in the main population simulations, most of the simulated afferent are not directly under the edge, we also compared the effect of moving the edge away from the RF center on the evoked response.

We found that the responses of simulated nerve fibers largely overlapped those of recorded ones, despite the fact that the former were not designed to mimic the latter, that is, were fit using the responses of different nerve fibers to different stimuli (**Error! Reference source not found**.). Indeed, measured spiking patterns, which were highly variable across nerve fibers, were as dissimilar to each other as they were to their simulated counterparts (**Error! Reference source not found.A-B**). Second, moving the edge around the RF had similar effects on measured and simulated afferent responses: Spike count decreased (**Error! Reference source not found.C**) and the latency of the first spike increased in a very similar fashion as the edge was moved away from the RF center (**Error! Reference source not found.D**).

#### Data analyses

We sought to test how precisely edge orientation can be decoded from the SA1 and RA population activity. Importantly, edge orientation was decoded while other parameters – indentation depth (and thus indentation velocity), aspect ratio, and location – were systematically varied. That is, we assessed the degree to which orientation could be robustly extracted across changes in other stimulus parameters. Otherwise, decoding becomes trivial because nerve fibers produce such highly repeatable and stimulus specific spiking responses that almost any feature can be extracted as long as other features are held constant.

**Spike distance metric**. A spike distance metric was used to gauge the dissimilarity of pairs of spike trains at different time scales (Victor and Purpura, 1997; Mackevicius et al., 2012; Weber et al., 2013; Saal et al., 2017). In brief, the distance between two spike trains was obtained by computing the lowest possible cost incurred to transform one spike train into another. Adding or deleting a spike incurred a cost of 1, whereas shifting a spike in time incurred a variable cost of q per unit time (measured in seconds). The temporal resolution of this metric could thus be adjusted by varying q: When q=0, the distance between two spike trains is solely determined by the difference in spike count; as q increases, shifting spikes becomes costlier than adding or removing them, and the timing of individual spikes increasingly determines the spike distance. The maximum cost is reached when removing all spikes in the first spike train and adding them back in the second one. Multiple q values were tested, ranging from 0 (equivalent to spike count) to 1000 (millisecond precision).

##### Decoding orientation using machine learning

*Input features*. For each afferent, we measured the spike distance between its responses to every possible pair of stimuli, yielding a 3750 x 3750 matrix. The resulting distance matrices were then combined across afferents (by taking the square root of the sum of squares) to obtain a distance matrix for the population response. Classical multidimensional scaling was then performed (CMDS, using cmdscale in MATLAB) to convert the population distance matrix to Euclidean coordinates, which were then used as input feature vectors to machine learning decoders. For q=0, coordinates are equivalent to the spike count vector; at non-zero values of q, distance is in part determined by spike timing, at a resolution that increases as *q* increases.

*Decoders*. We implemented three different decoding methods to estimate edge orientation: Linear regression (LIN, using fitrlinear in MATLAB), linear discriminant analysis (LDA, using fitcdiscr in MATLAB), and a simple feedforward artificial neural network (ANN, using fitnet in MATLAB), with a single hidden layer consisting of 10 units (as more nodes or layers did not yield better performance). The goal of this analysis was to estimate the upper bound on achievable performance and the decoder complexity necessary to achieve it (ranging from linear regression to ANN). To characterize the time course over which orientation information conveyed by the afferent population evolves, classification analyses were performed over progressively expanding time windows, ranging in duration from 1 to 500 ms and spaced logarithmically. A new set of parameters was fit for each time window. Decoding performance was evaluated using ten-fold cross-validation. That is, the data were randomly split into 10 equally sized sets, one of which was retained as the validation set to test the performance of the decoder and the other nine were used as training sets for the fitting procedure. This process was repeated 10 times, with each of the 10 sets used as validation set. In the case of the ANN, for each fold of the fitting procedure, the training sets were further split into two subsets, one for training and one for testing (90/10 percent of the data, respectively). As high-dimensionality in the input space can lead to overfitting, we reduced the number of dimensions of the input feature vector before training by selecting a restricted number of principal components (PCs) resulting from the CMDS. Under most circumstances, best cross-validated performance was achieved with the first 100 PCs.

**Decoding orientation using a biologically inspired decoder**. As mentioned above, the machine learning decoders gauge the degree to which orientation information is contained in the responses of populations of afferents, without consideration of how these responses might be interpreted downstream. The performance of these decoders thus constitutes an upper limit on the informativeness of afferent signals about orientation. In light of this, we wished to assess whether orientation could be decoded from afferent responses using a biologically inspired mechanism. As briefly described above, edge orientation has been hypothesized to be extracted from the spatial pattern of activation across nerve fibers by implementing a spatial filter in somatosensory cortex that consists of excitatory regions flanked by inhibitory ones, a RF structure well approximated by a Gabor filter (DiCarlo and Johnson, 2000; Bensmaia et al., 2008a). With this in mind, we built a “biomimetic” decoder based on the same principle: A population of downstream neurons (“cortical neurons”) was sampled on a square grid centered on and blanketing the stimulus (14 mm wide, spaced 0.35 mm apart, yielding 41×41 neurons), each described as a Gabor filter. While Gabor filters oversimplify the response properties of cortical neurons – for example by omitting the lagged inhibitory component (DiCarlo and Johnson, 2000) –, they reliable predict cortical responses to edges (Bensmaia et al., 2008b). For each cortical neuron, the cumulative spike counts of all afferents were weighted according to the respective positions of their RFs within a spatial Gabor function, representing the cortical neuron’s RF, then summed to determine the cortical neuron’s response. The Gabor filter is a 2D symmetric filter, f(x, y) of the form:

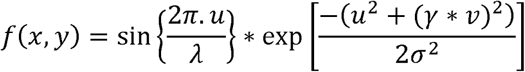

with *u = (x − x_c_) * cos(θ) + (y − y_c_) * sin(θ), v = −(x − x_c_) * sin(θ) + (y − y_c_) * cos(θ)*

where (x, y) denotes the spatial position of the tactile fiber’s RF, (*x_c_, y_c_*) denotes the center of the filter, *θ* represents the orientation of the Gabor, *σ* is the standard deviation (or width) of the 2D Gaussian, *λ* is the wave length and *γ* is the aspect ratio. Multiple populations of cortical neurons were simulated, each with a different orientation, *θ*, to mimic the response of different cortical subpopulations, each selective to a different orientation. For simplicity, the preferred orientations of the various subpopulations were aligned to the stimulus orientations. Finally, the responses of all cortical neurons in each subpopulation were rectified and summed to yield a scalar value. The decoded orientation was that of the population yielding the maximum value (i.e., winner-take-all). Gabor parameters *σ*, *λ*, and *γ* were optimized to minimize the root mean square error of the orientation prediction over the stimulus set, and constrained over the range of biologically plausible values (*σ* = 3.1, *λ* = 7, expressed in mm, and *γ* = 1.6 cf. (DiCarlo and Johnson, 2000)).

**Decoding orientation using first spike recruitment sequence**. In analyzing afferent responses with spike distance metrics as described above, we assess the degree to which the timing of spikes emitted by individual afferents carries information about orientation. This analysis does not account for the possibility that the relative timing of responses across afferents carries stimulus information. For example, the sequence in which afferents are recruited by a tactile stimulus has been put forth as a potential code for feature coding in touch (cf. Johansson and Birznieks, 2004). Indeed, the order in which tactile fibers are activated depends on the spatial configuration of the tactile stimulus. Furthermore, this recruitment sequence can be decoded downstream and is, to some extent, robust to noise (Bichler et al., 2012). In light of this, we implemented the recruitment sequence decoder on our simulated responses. Since this mechanism has been shown to extract tactile features very rapidly, we used it as a benchmark with which to compare the time course of the biomimetic spatial decoder. Specifically, afferents were sorted by the latency of their first spike for each trial (out of a total of 3750 trials). The mean recruitment sequence was then obtained for all condition (125 different conditions: 5 depth x 5 length x 5 repetitions) for each of the 30 orientations. Finally, each single-trial recruitment sequence was correlated (using Spearman’s rank correlation coefficient) with the mean recruitment sequence for each orientation excluding that trial (i.e., using a leave-one-out procedure). The orientation yielding the highest correlation was taken to be the decoded orientation. Decoding error was defined as the angular difference between the decoded orientation and the actual orientation. This decoding procedure is almost identical to a previously implemented one (Johansson and Birznieks, 2004). To assess the evolution of decoding performance with time, the procedure was repeated for 30 progressively expanding time windows, ranging from 1 ms to 500 ms after stimulus onset and spaced logarithmically (as for previous analyses).

### Responses of neurons in somatosensory cortex

The biologically plausible decoder is inspired by known response properties of cells in somatosensory cortex. The analysis with the biologically plausible decoder establishes a time course at which we would expect orientation signals to evolve in cortex. To validate this predicted time course, we analyzed the responses of single neurons in the somatosensory cortex of non-human primates. As these data have been previously published (Bensmaia et al., 2008a), the experimental methods are only summarized here.

#### Experimental procedures

**Stimuli**. The stimulator used was the same as in the peripheral recordings described above, and most experimental parameters were identical (edge length and width, indentation depth, orientations, and ramp durations). On each trial, a bar was indented into the skin and held for 100 ms. The pivot of the bar was always located at the point of maximum sensitivity (or hotspot) of the neuron. The inter-stimulus interval was 100 ms. Edges were each presented 10 times in pseudorandom order for a total of 80 trials (8 orientations x 10 presentations).

**Neurophysiology**. Extracellular recordings were made in the postcentral gyri in one hemisphere of each of two awake Rhesus macaques (Macaca mulatta, 1 female). Recordings were obtained from neurons in areas 3b and 1 whose RFs were located on the distal pads of digits 2–5, and that met the following criteria: (1) action potentials were well isolated from the background noise, (2) the RF of the neuron included at least one of the distal finger pads on digits 2–5, (3) the stimulator array could be positioned so that the RF of the neuron was centered on the array, and (4) the neuron was clearly driven by the stimulus.

#### Data analysis

**Single neurons**. For each neuron, we characterized the time course over which the orientation signal emerged in its spiking response. The strength of the orientation signal was evaluated by estimating the discriminability of the single unit responses to two edges at two orientation, using a d’, as described in (Berens et al., 2012),

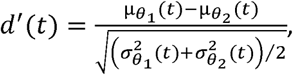

where µ and σ are the mean and standard deviation of the firing rate across repetitions. Smooth time-varying firing rates were obtained by binning spike trains in 1 ms bins, and then filtering the binned vectors with a trailing moving average of 25 bins. This analysis was carried for each pair of orientations and each neuron.

**Population**. The same cross-validated decoding procedure as described for the peripheral afferent simulation was used for the cortical population. The input vector of the decoder was the spike count vector (q=0) reduced to its first 10 PCs. The decoding window, starting from stimulus onset, expanded to 100 ms in thirty equally spaced time steps. Given the similarity in performance of the three machine learning decoders, only the LDA algorithm was tested here. Since cortical neurons were not recorded simultaneously, we built in noise correlation using the following procedure. First, for each neuron and stimulus, the repetitions of each stimulus were sorted by spike count, leading to a high level of noise correlations (mean correlation = 0.8). Then, random pairs of repetitions were swapped for each neuron and each repetition until the mean noise correlation was 0.25, a value commonly observed in early cortical areas (Cohen and Kohn, 2011).

**Comparison with the periphery**. We simulated the responses of the afferent population to the stimuli used in the cortical experiment. We then compared the performance and time course of the biomimetic decoder based on afferent responses (using the same parameters as before) to their counterparts derived from cortical responses.

## Results

### Simulated population responses to indented edges

We simulated the spiking response of all SA1 and RA afferents that innervate the skin to edges indented into the fingerpad at different orientations, with different aspect ratios, to different depths, and at different locations (**Figure 2 A-C**). The simulation has been shown to reproduce with millisecond precision the responses of tactile nerve fibers across a wide range of conditions (Saal et al., 2017)(**Figure 1**). First, we examined the extent to which information about edge orientation could be extracted from the SA1 and RA population activity, irrespective of the other stimulus parameters (aspect ratio, depth, location), using a variety of standard machine learning classifiers. Second, we determined whether information about orientation was carried only in the spatial patterns of activation (in the spike counts) or if precise spike timing also contributes to the orientation signal. Finally, we investigated whether orientation information could be extracted using a decoder inspired by known response properties of downstream somatosensory neurons.

**Figure 1.**
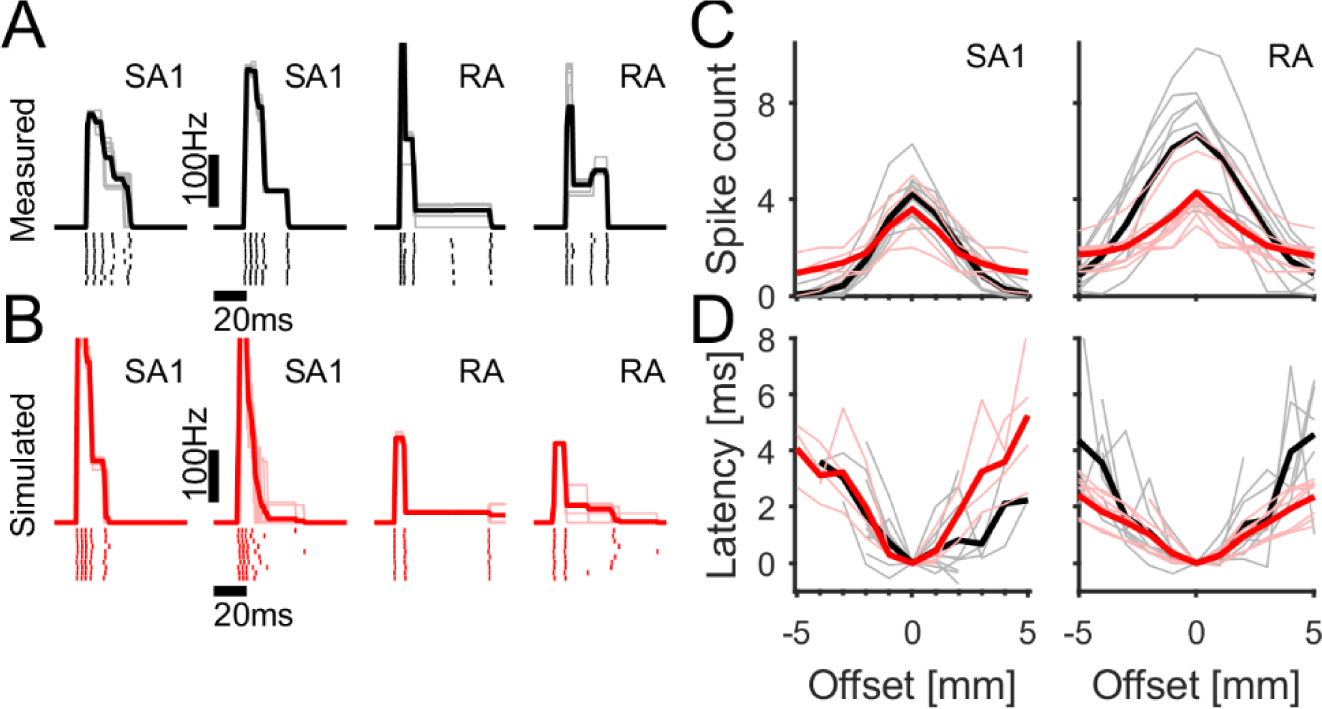
Simulations validation. **A-B|** Comparison of measured (A, in black) and simulated (B, in red) single unit response to bars indented in the center of their RFs. Typical responses of two SA1 and two RA fibers. The rasters of the responses evoked by 10 repetitions are shown together with their firing rate profiles as a function of time. **C-D|** Number of spikes (C) and latency of the first spike relative to the center of the RF (D) evoked by an indentation as a function of distance of the bar from the center of the receptive field. Black are recorded and red are simulated data. Thin light line denote means across repetitions from single units, thick dark traces are averages across units. Simulated responses are overlap with their measured counterparts even though the models were not built to mimic these particular nerve fibers.

As expected, SA1 fibers responded strongly at the onset of the stimuli, and some of them maintained their responses throughout the hold phase, whereas the RA fibers only responded during the onset and offset ramps (**Figure 2 D-E**). Each stimulus activated a large fraction of the afferents innervating the fingerpad and most afferents responded to at least one stimulus, as observed previously (Bisley et al., 2000; Jenmalm et al., 2003). Visual examination of the spatial layout of SA1 and RA population responses suggests that it reflects the orientation of the indented edge. The spatial layout of the edge was reflected in the spatial pattern of activation in both SA1 and RA responses, though SA1 fibers conveyed a more acute spatial image, as expected (**Figure 2 D-E, right**).

**Figure 2.**
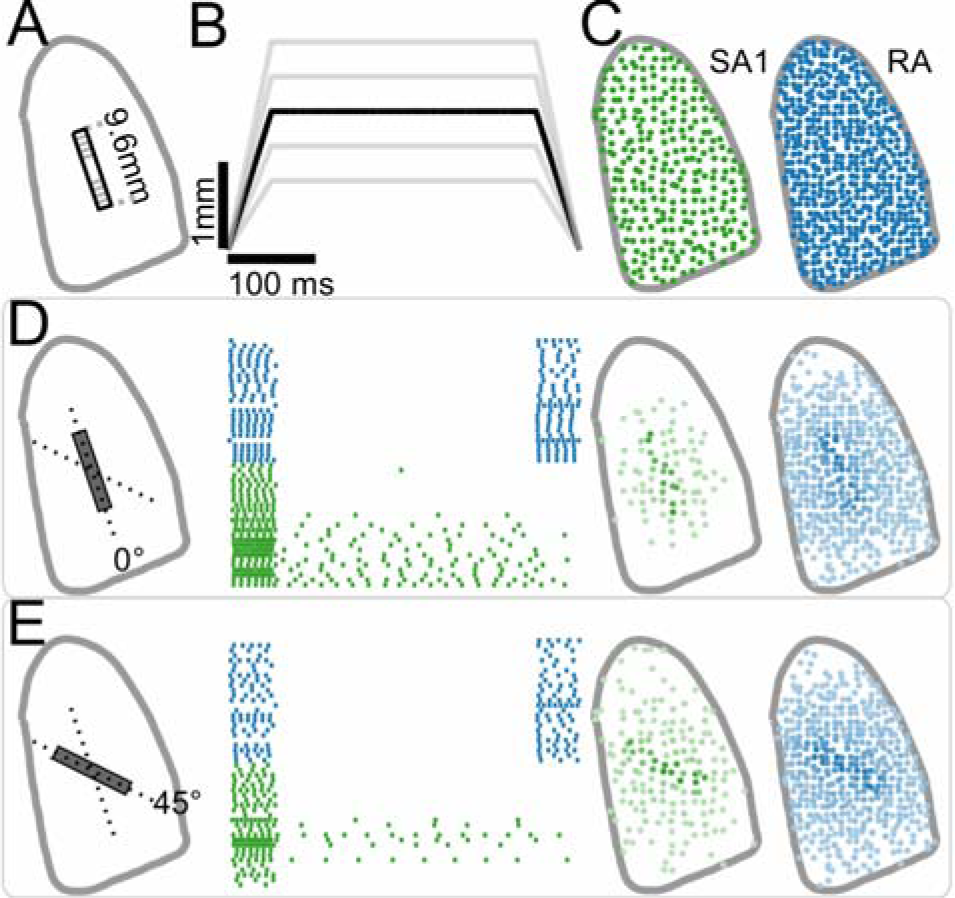
Simulated population response of tactile fibers to edges indented into the skin. **A|** Rectangular bars of different lengths (ranging from 3.2 to 9.6 mm) and orientations (ranging from 0 (A, D) to 45 degrees (E)) are indented into the skin. **B|** Indentation trajectory at different indentation depths (ranging from 0.8 to 2.4 mm). **C|** SA1 (left, green) and RA (right, blue) afferents tile the fingerpad. **D-E|** Population response to two indented bars (length=9.6 mm, depth=2.4 mm) at different orientations: 0 degrees (D), and 45 degrees (E). Center: raster of the 25 most active tactile fibers of each type across all directions. Right: Spatial distribution of the strength of the response, darker points denote more active afferents.

While changes in orientation did not have a systematic effect on population firing rates, changes in other stimulus features did (**Figure 3**). Indeed, increases in bar length and indentation depth both led to an increase in the aggregate firing rate (**Figure 3 A-C**). Changes in indentation depth also affected the timing of the population response as gauged with the first and second spike latencies (**Figure 3 D**). Variations in these other stimulus features are thus liable to obscure orientation signals.

**Figure 3.**
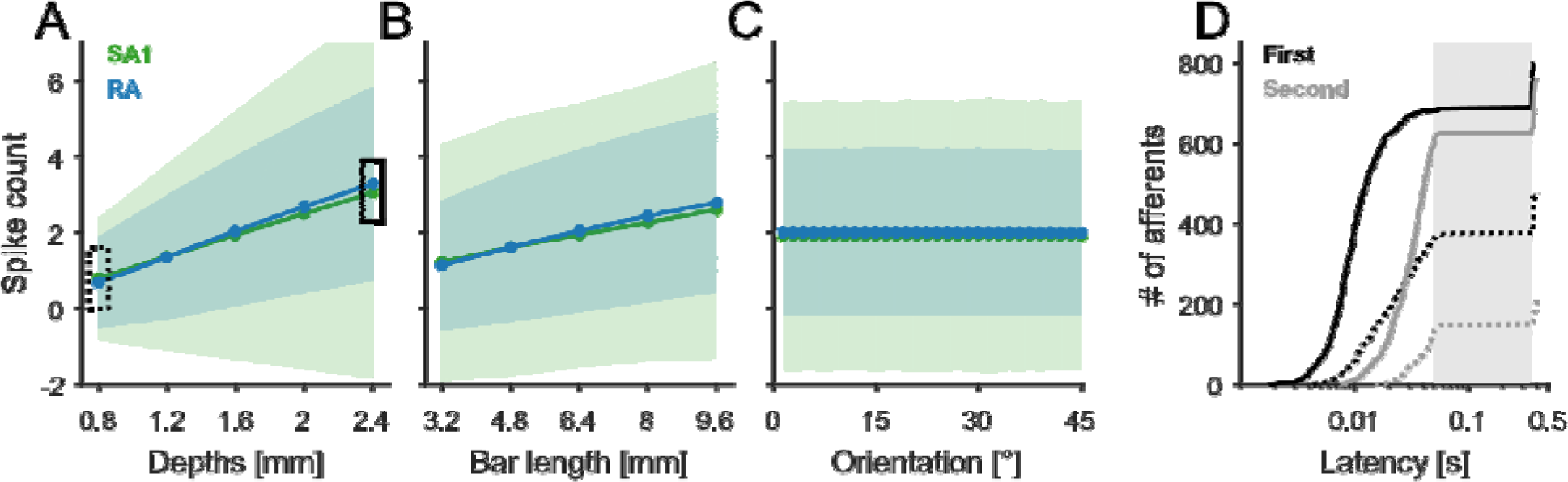
Sensitivity of the population response to stimulus parameters. **A, B,C|** Mean evoked spike count as a function of edge depth (A), length (B) and orientation (C). Shaded areas show one standard deviation. **D|** Cumulative distribution of the first (black) and second (gray) spikes latency for two trials at different indentation depths (0.8 mm, dotted lines; 2.4 mm, solid lines) Orientation was 1.5° and bar length was 9.6 mm. Grey shaded area indicates the hold period. Responses to the same stimulus vary widely across nerve fibers.

### Orientation signals in population responses

First, we wished to assess the degree to which orientation could be decoded from spatio-temporal patterns of afferent activation without regard to biological implementation (**Figure 4**). To this end, we trained three decoders of increasing complexity to classify edge orientation using the collective spiking response of the afferent population. Initially, we tested decoders based on spike counts alone. We found that decoders achieved high classification accuracy (∼ 5° error) within 10 ms (**Figure 4 A**) – when most afferents had only fired one spike –, and leveled off at 1-3° within 50 ms. While the simple linear decoder yielded high accuracy, the more elaborate ones, including LDA and ANN, were even better. Decoding performance was better for deeper indentations (**Figure 4 B**), as might be expected given the effect of depth on spike count (**Figure 3 A**), and, of course, for longer edges (**Figure 4 C**), as has been shown for perceptual angular acuity. Finally, jittering stimulus position led to poorer orientation performance for all decoders (**Figure 4 D**).

**Figure 4.**
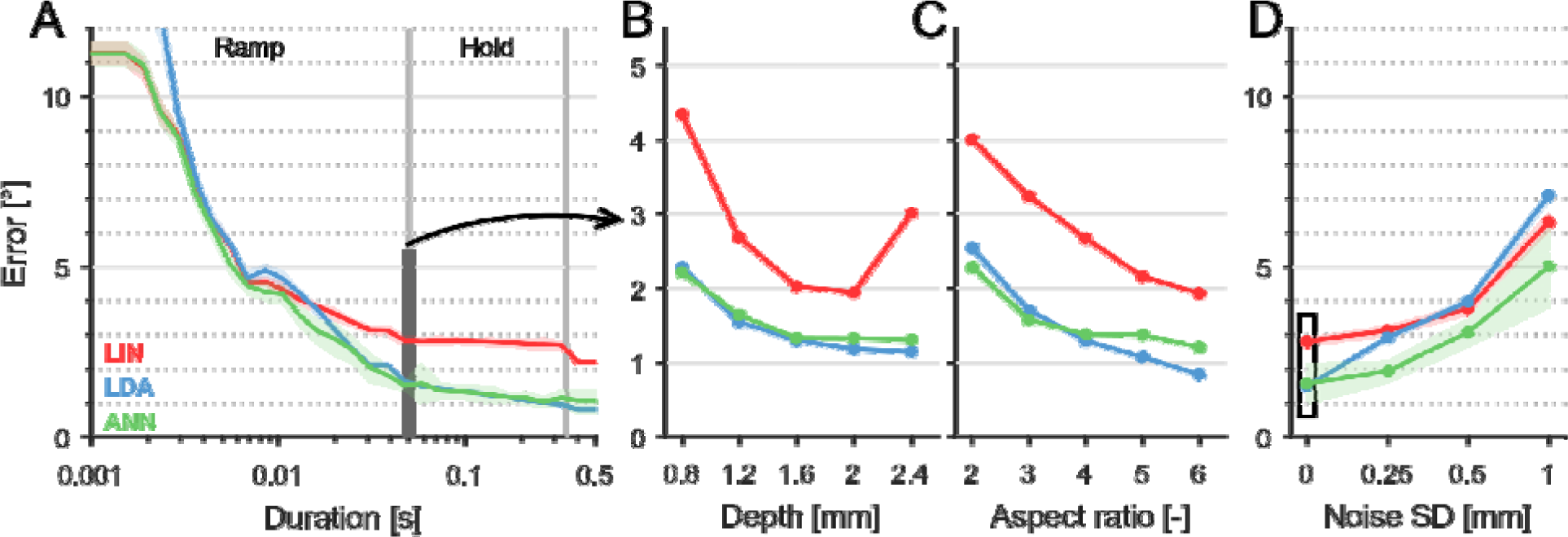
Orientation decoding performance based on spike count (q=0). **A|** Evolution of decoding performance as a function of stimulus duration after stimulus onset (with zero noise). **B, C|** Mean decoding performance 50 ms after stimulus onset (indicated by a vertical black line in panel A), as a function of indentation depth (B) and length (C). As expected, performance improves as indentation depth and edge length increase. **D|** Orientation decoding performance as a function of noise level. The black box corresponds to the data shown in panel A. Different colors denote different decoding approaches. In A and D, lines and shaded areas denote mean and std across the 10-fold cross-validation, respectively (some std’s are hardly visible because they are diminutive).

Next, we examined whether taking into consideration precise spike timing improves decoding performance since orientation decoding performance based on the spiking responses of individual nerve fibers has been found to peak at a temporal resolution of about 2-4 ms (Mackevicius et al., 2012; Weber et al., 2013; Pruszynski and Johansson, 2014; Suresh et al., 2016). When decoding orientation from population responses, however, taking spike timing into account did not have a major impact on decoding performance. Indeed, only in the early response (before 10-20 ms) in the noise-free condition was a slight improvement in performance observed (**Figure 5 A**). However, after that period and in all noisy conditions, performance did not improve with the inclusion of spike timing and actually worsened during the hold phase or at high temporal resolutions (**Figure 5 B**). At 50 ms after onset, LIN and LDA actually performed slightly better with temporal resolutions around 20 ms (q’s between 30 and 100 s^−1^), but this performance improvement disappeared for longer time windows and at high levels of noise.

**Figure 5.**
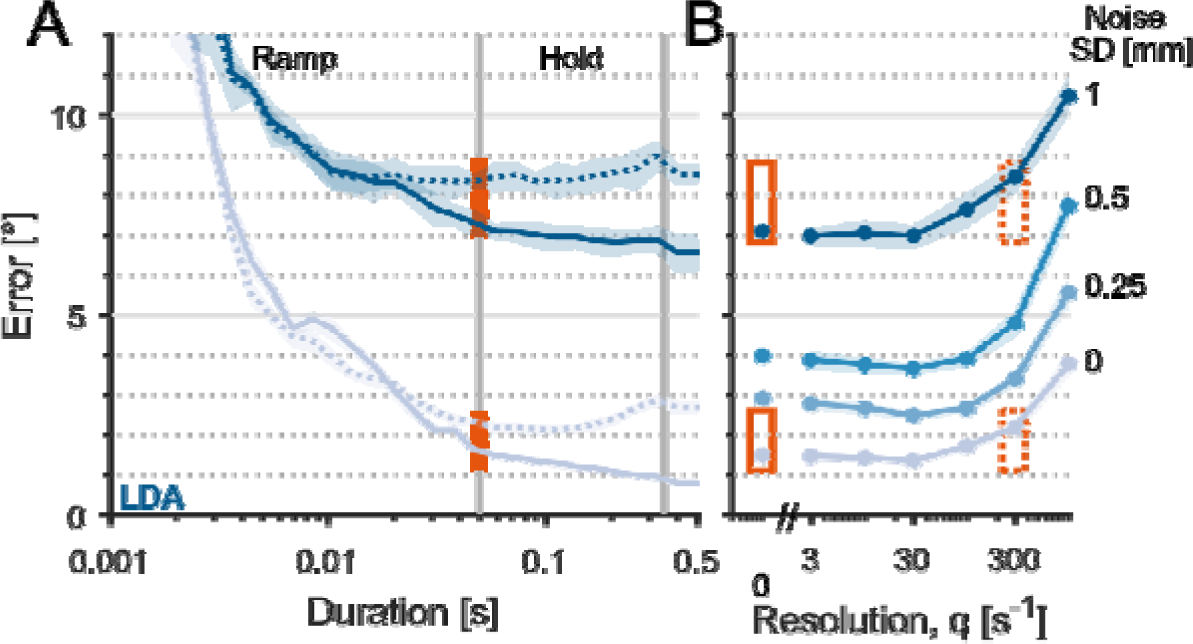
Taking spike timing into account has little impact on decoding performance. **A|** Evolution of decoding performance as a function of stimulus duration after stimulus onset for the LDA algorithm at two extreme noise levels (SD=0 and 1 mm). Solid lines denote performance based on spike count (q=0), dotted lines performance with spike timing (q=300). **B|** Average decoding performance 50 ms after stimulus onset (depicted by the orange box in panel A), as a function of q, the temporal resolution of the spike distance metric (see methods). The four levels of noise are shown (SD = 0, 0.25, 0.5 and 1mm, from light to dark) Boxes depict the examples shown in A. Lines and shaded areas show cross-validated mean ± std error.

Finally, we examined the orientation signals conveyed by each class of afferents separately (SA1 and RA). The performance levels for each sub-population were slightly poorer than that for the entire population (**Figure 6**) and similar to each other, as observed previously (Saal et al., 2017). RA-mediated orientation signals tended to be faster than their SA1 counterparts, but the SA1-mediated signal continued to improve during the hold phase, when RA fibers are silent. Precise spike timing did not provide substantial performance improvement for within-population decoding (**Figure 6**, dashed lines).

**Figure 6.**
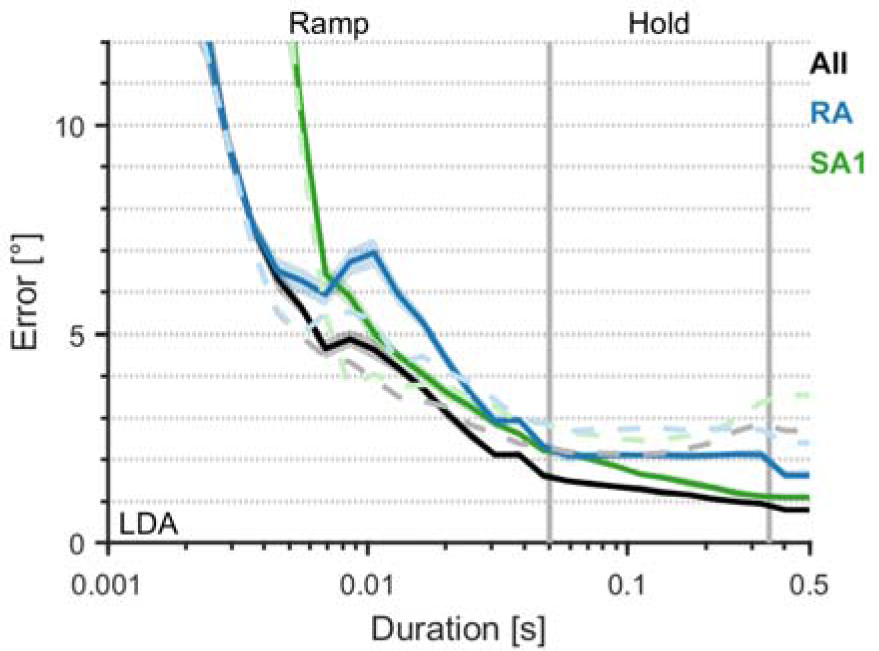
Performance of different tactile submodalities. Evolution of decoding performance as a function of stimulus duration after stimulus onset for the rate-based LDA (zero noise). Black line shows the performance for all afferents, green lines show the performance based only on SA1 responses and blue line show performance based on RA responses. The dashed light lines show performance with precise spike timing included (q=300). Lines and shaded areas show cross-validated mean ± std error.

In summary, the spatial pattern of activation across afferents rapidly conveys accurate information about edge orientation across a wide range of indentation depth and lengths, and precise spike timing does not convey much additional information.

### Orientation decoding using a biologically plausible mechanism

With the previous analyses, we showed that orientation could in principle be decoded quickly and accurately from spatial patterns of activation across afferent populations. That orientation information is carried in these signals does not guarantee that this information is or even can be extracted by a neural circuit. With this in mind, we examined whether orientation could be classified using a biologically inspired decoder. Specifically, the RFs of neurons in somatosensory cortex have been shown to comprise excitatory regions flanked by inhibitory ones and this RF structure, well approximated using a Gabor function, accounts for the neurons’ orientation tuning (DiCarlo and Johnson, 2000; Bensmaia et al., 2008a). With this in mind, we convolved the spatial layout of the response, expressed as a spike count, evoked in each fiber by each stimulus with Gabor filters at different orientations. The orientation that yielded the highest value of this convolution was then selected (see Methods).

First, we estimated the degree to which edge orientation could be decoded using the biomimetic decoder and characterized the time course over which performance evolved in the absence of jitter in the stimulus position (**Figure 7 A**). While discrimination accuracy was poorer for the biomimetic decoder than for machine learning algorithms, as expected, its performance was still very good, leveling off at an acuity of around 3-4° after only 50 ms. The dip in performance around 10 ms observed in **Figure 7 A** (and also seen to a lesser degree in **Figure 4 A**) is due to a second wave of spikes, which temporarily obscures the orientation (see **Figure 3 D**). Second, we investigated the impact of aspect ratio on orientation decoding performance and found that, as expected, performance improved as aspect ratio increased (**Figure 7 B**). Importantly, the performance of the biomimetic decoder greatly surpassed that of human observers performing a corresponding angular discrimination task (Pruszynski et al., 2018). Moreover, the biomimetic decoder was far less affected by jitter in the position of the stimulus than were the machine learning decoders. Indeed, performance decreased only slightly as the position jitter increased.

**Figure 7.**
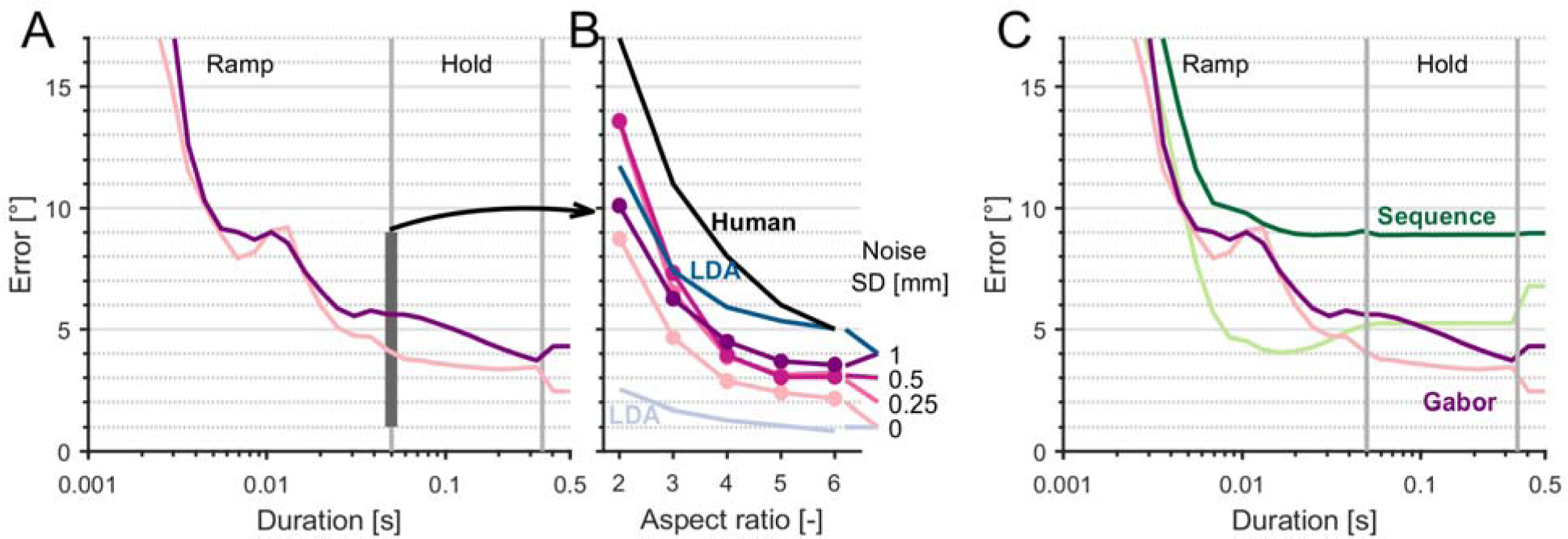
Biomimetic decoder also resolves orientation quickly and accurately. **A|** Evolution of decoding performance of the biomimetic decoder as a function of stimulus duration after stimulus onset at two extreme noise levels (SD=0 and 1 mm). **B|** Decoding performance 50 ms after stimulus onset (purple, denoted by the thick vertical line in A) as a function of aspect ratio, compared to human performance (black, adapted from (Pruszynski et al., 2018)) and to ML performance (blue dashed line, same as in figure 2C). The four levels of noise are shown for the biomimetic decoder (SD = 0, 0.25, 0.5 and 1mm, from bright to dark), the two extreme values are shown for the LDA decoder. **C|** Performance of the biomimetic decoder (purple traces, also shown in panel A) compared to a decoder based on recruitment sequence (green traces).

We then compared the performance of the biomimetic decoder to a decoder using the temporal sequence of recruitment of the first spike in all tactile fibers, a neural code that has been shown to carry information about complex spatial fingertip events shortly after stimulus onset (Johansson and Birznieks, 2004). We found that the recruitment sequence signaled orientation with an acuity below 5° within 10 ms in the low noise condition (**Figure 7 C**), 20 ms faster than the biomimetic decoder. However, like the machine learning decoders, the recruitment sequence decoder was far less robust to noise, peaking in acuity at 9 degrees when a high level of noise was introduced.

Thus, while the spike count-based biomimetic spatial decoder is somewhat slower than the spike timing-based recruitment sequence decoder, the biomimetic decoder is still very rapid and more robust to noise.

### Orientation signals in cortex

As discussed above, the biomimetic decoder – a spatial Gabor filter – is designed to mimic the computation that is implemented by neuronal circuits to culminate in orientation tuned responses in somatosensory cortex (DiCarlo et al., 1998). With this in mind, we wished to characterize the time course of orientation signals in cortex and compare them to those extracted by the biomimetic decoder. We expected the former to mirror the latter after accounting for transmission delays. To test this hypothesis, we re-analyzed responses of neurons in somatosensory cortex – including Brodmann’s areas 3b and 1 – to edges indented into the skin (using data previously published in (Bensmaia et al., 2008a)).

The spiking activity of 163 neurons in somatosensory cortex (52 in area 3b and 111 in area 1) was recorded when edges were rapidly indented into the skin at the center of each neuron’s RF at a depth of 500 µm, held for 100 ms, and then rapidly retracted (**Figure 8**, stimulus on-and off-ramps depicted by blue shaded areas). As described previously, a large fraction of cortical neurons are orientation selective (**Figure 8 A**): they respond most strongly to an edge at a given orientation and their response drops off smoothly as the orientation deviates from this preferred orientation. This orientation tuning can be accurately accounted for by a model that describes neuronal RFs as Gabor filters (DiCarlo et al., 1998; Bensmaia et al., 2008a).

**Figure 8.**
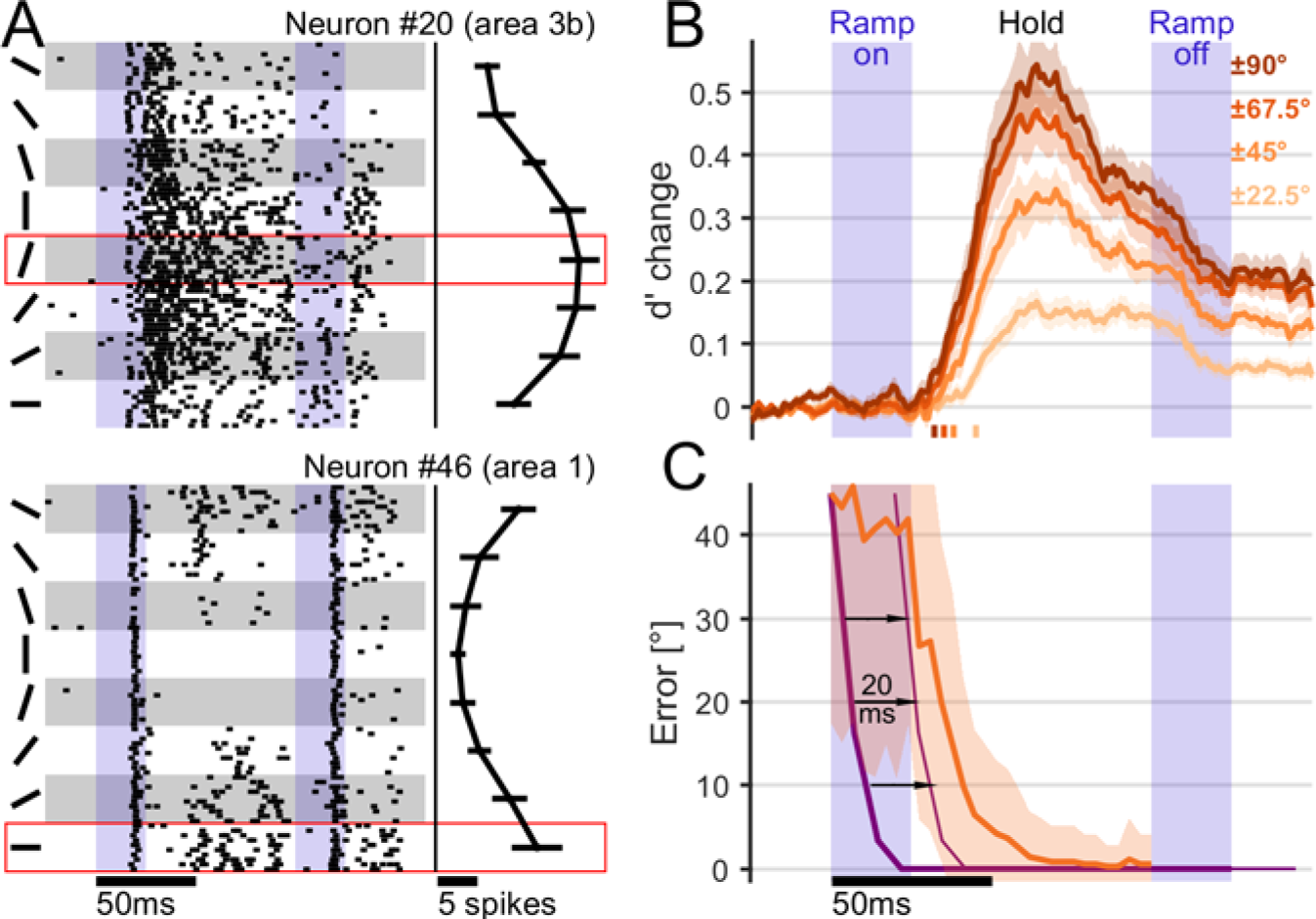
Evolution of orientation signals in somatosensory cortex. **A|** Response of two typical orientation sensitive neurons in areas 3b and 1 to indented edges. Left: Raster plot of the response to 10 repeated presentations of edges at 8 orientations ranging for 0° to 180°. Right: Mean (±std) spike count over the entire trial. Each neuron’s preferred orientation is indicated by the red box. The blue shaded areas depict stimulus onset and offset ramps. **B|** Evolution of d’ in individual cortical neurons as a function of stimulus duration. The solid lines show the mean across all neurons (n=197), the shaded area shows ±1 sem. Different colors denote different pairs of angles (±22.5°, ±45°, ±67.5° and ±90°). Tick marks denote the time at which d’ becomes significantly different from baseline (t-test, p<0.01) **C|** Evolution of orientation decoding performance from the cortical population (n=197, reduced to 10 PCs in orange) as a function of stimulus duration, using LDA. For comparison, orientation decoding performance from the afferent population using the biomimetic decoder (purple, thick line). The same data are reproduced with a lag of 20 ms (thin line) to account for the conduction delay to cortex. Lines and shaded areas show mean ± std across the 10-fold subsets of validation data.

First, we examined the evolution of orientation selectivity in single-unit responses as gauged by the discriminability of pairs of edges at different orientations using d’ (see Methods). We found that d’s from single unit responses begin to ramp up as soon as the stimulus-evoked response appears. A significant change from baseline was observed as early as 37.5 ms after stimulus onset on average, so about 17.5 ms after response onset (**Figure 8 B**, p < 0.01). The d’ peaked around 60 ms after stimulus onset, following a time course consistent with orientation signals in afferent populations after accounting for conduction delays (∼ 20 ms). At the population level, we observed a slightly faster time course, with decoding error dropping below 2° within 60 ms (**Figure 8 C**). After incorporating the conduction delay, decoding performance based on afferent and cortical responses followed very similar time courses. The performance of the decoder from cortical responses was not influenced by the level of noise correlation (see Methods), as found previously in V1 (Berens et al., 2012).

In conclusion, orientation signals emerge very rapidly in cortex and with a time course coherent with that of orientation signals in the nerve.

## Discussion

### Limitations of the simulation

The present results hinge on analyzing the responses of all nerve fibers activated by the stimulus and thus could not be achieved without the nerve simulation. Indeed, measuring the response of the entire afferent population is currently impossible as is reconstructing the population response from the sequentially recorded responses of individual fibers given the dependence of afferent responses on the position of their RF with respect to the stimulus. While the model provides a precise and accurate simulation of afferent responses (Phillips et al., 1981; Sripati et al., 2006; Saal et al., 2017), it does not capture some of the variation in sensitivity and RF shape that is observed in the nerve (Johansson, 1978; Goodwin et al., 1995; Pruszynski and Johansson, 2014). However, these deviations from the true population response are unlikely to substantially affect the signaling of geometric features if these are decoded spatially as has been shown in reconstructions of the time-averaged spatial layout of the afferent responses to such features (Goodwin and Wheat, 2002).

Another possibility is that the model mis-or under-estimates trial-to-trial variability in afferent responses, which may inflate the orientation decoding performance. However, our manipulation of stimulus jitter shows that the biomimetic orientation decoder is relatively impervious to noise.

Finally, the model simplifies the mechanics of the skin and its geometry (Saal et al., 2017). For example, the curvature of the finger is not included in the model, nor is the presence of bone, which may be particularly relevant near finger joints, where the bone is most superficial. However, most object interactions involve the fingerpads (Christel et al., 1998), so will be well approximated by the model. Whether the curvature of the finger and the underlying bone have systematic effects on feature signaling remains to be determined.

### Comparison with previous approaches

Previous attempts to understand how spatial patterns are represented in the responses of afferent populations have adopted one of two approaches. In the first approach, the stimulus is moved relative to the RF of a given fiber, and the population response is reconstructed by tiling the responses of the afferent over the stimulus, yielding a so-called spatial even plot (Phillips and Johnson, 1981; Goodwin et al., 1995; Wheat et al., 1995). The advantage of this approach is that it systematically tiles the skin (though typically not accounting for the density of innervation); the disadvantage is that variability across nerve fibers and the cutaneous innervation density are generally not taken into account (but see (Goodwin and Wheat, 2002)) and only one class of afferents is considered at a time. In the second approach, the stimulus is presented at a single location, typically the center of the distal fingerpad, and responses are collected (non-simultaneously) from afferents with RFs at different locations on the fingertip (Khalsa et al., 1998; Bisley et al., 2000; Birznieks et al., 2001, 2010; Friedman et al., 2002; Jenmalm et al., 2003). The advantage of this approach is that it preserves the idiosyncratic properties of different afferents but provides a sparse representation of the full neural response.

While both approaches converged on the conclusion that SAI and, to a lesser extent, RA fibers, provide a spatial image of the stimulus that could in principle signal the spatial features of the stimulus (Goodwin and Wheat, 2004), responses were described in terms of total spike count or mean firing rate, thereby obscuring the time course at which these representations emerge. Furthermore, because these representations of the neuronal response are incomplete, they do not provide a reliable estimate of the true acuity of the afferent signal.

Moreover, the impact of varying orthogonal stimulus parameters, such as indentation depth or position, was not assessed, so the robustness of these representations could not be evaluated. For example, geometric features, such as curvature, can be extracted from the relative timing of the first or second spike of the population response when other stimulus parameters are held constant (Johansson and Birznieks, 2004). The question remains, however, whether such a mechanism can decode surface curvature across changes in force direction, for example. Indeed, while single afferents convey orientation signals in the timing of their responses (Pruszynski and Johansson, 2014), such signals are highly dependent on other stimulus features (Suresh et al., 2016). Our results suggest that signals that rely on precise spike timing, at least using the two decoders we implemented, are especially susceptible to small changes in stimulus delivery (**Figure 3D**), which renders this signal about geometric features much more ambiguous under naturalistic conditions. We do not claim there exists no other mechanism that could extract information about geometric features from precise spike timing, for example at the level of populations of nerve fibers through a coincidence detection mechanism (Pruszynski and Johansson, 2014). Neither do we claim that spike timing plays no role in tactile coding, as millisecond level spike-timing of afferent responses has been shown to play a role in the coding of texture (Weber et al., 2013) and of the frequency of skin vibrations (Mackevicius et al., 2012; Birznieks and Vickery, 2017). The present results demonstrate, however, that the classical model of orientation signaling in touch – broadly analogous to its visual counterpart – operates on a time scale that is rapid (and precise) enough to guide object interactions without taking spike timing into account.

### Tactile hyper acuity

We show that edge orientation can be decoded with a precision of 2 degrees within 50 ms (for long edges). A rotation of 2 degrees corresponds to a displacement of only 0.16 mm at edge tips, far smaller than the inter-afferent spacing (∼0.7/mm at the fingertip), thereby constituting an instance of tactile hyper-acuity (Loomis, 1980). How then is it possible to extract orientation so accurately? While the interplay of RF hotspots and a mechanism of coincidence detection has been proposed as a mechanism to achieve resolutions beyond afferent density and explain human acuity (Lechelt, 1992; Dodson et al., 1998; Bensmaia et al., 2008a; Peters et al., 2015; Pruszynski et al., 2018), we show that hyperacuity can arise from spatially patterned, graded activity of adjacent afferents with overlapping RFs. The limits to tactile acuity are not set just by the innervation density, but also by the resolution of the graded activity, the integration time window, and the noise. In fact, that the orientation acuity of a single afferent population nearly matches that of the entire population suggests that innervation density is actually not the limiting factor, contrary to the consensus view (Goodwin and Wheat, 2004).

### Comparison with other sensory areas

The responses of single neurons in somatosensory cortex become rapidly tuned to orientation (within 40 ms with a peak around 60 ms, **Figure 8 B**), leading to a rapid extraction of the orientation signal at the level of the population (within 60ms, **Figure 8 C**). Similar results have been reported in visual cortex (V1), where orientation signals are present at response onset (80 ms after stimulus onset)(Berens et al., 2012) and peak 40 ms later. Orientation information emerges more rapidly in somatosensory cortex due to the shorter response latencies. While a circuit model of the pathway that gives rise to orientation tuning has been proposed and substantiated (Hubel and Wiesel, 1962; Clay Reid et al., 1995), no such model yet exists for the sense of touch, due to the paucity of studies investigating the RF properties of neurons residing in intermediate stages of somatosensory processing, namely the dorsal column nuclei and thalamus.

### Conclusions

A wealth of information about objects and about our interactions with them is carried in neural signals from the glabrous skin of the hand. As information is multiplexed in these afferent responses, the read out of this information by downstream neurons depends on the relevant stimulus property, each property mediated by its own code(s). For example, the magnitude of a tactile stimulus is encoded in population-level spike count (Muniak et al., 2007; Graczyk et al., 2016), fine surface texture or vibratory frequency is encoded in precise spiking patterns (Saal et al., 2016), and, according to the classical view, geometric features are encoded in spatial patterns of activation across the sensory sheet (Goodwin and Wheat, 2004).

In this study, we show that information about edge orientation – encoded in spatial patterns of afferent activation – emerges rapidly, starting with the very first spike, challenging the intuition that such this spatial code would be slow because it requires integrating spikes over time. Furthermore, these spatial signals can be decoded reliably using a biologically plausible mechanism that matches the properties of neurons in somatosensory cortex. Orientation signals carried by these neurons, in fact, follow a time course that is entirely consistent with the proposed decoding mechanism. Finally, we show that the spatial representation can account for hyper acuity: Measured angular acuity in humans exceeds that naively predicted from the cutaneous innervation density but is predicted precisely by the proposed orientation processing mechanism. These results illustrate how a complete population may carry information in a way that cannot be inferred from the responses of individual neurons or even subpopulations of neurons.

## Acknowledgments

We thank Peter Denchev for assistance with the cortical data collection and Hannes Saal and Justin Lieber for useful comments on a previous version of the manuscript. This work was supported by NINDS grants NS101325 and NS095162.

## References

Bensmaia SJ, Denchev PV., Dammann JF, Craig JC, Hsiao SS (2008a) The Representation of Stimulus Orientation in the Early Stages of Somatosensory Processing. J Neurosci 28: 776–786.

Bensmaia SJ, Hsiao SS, Denchev PV, Killebrew JH, Craig JC (2008b) The tactile perception of stimulus orientation. Somatosens Mot Res 25: 49–59.

Berens P, Ecker AS, Cotton RJ, Ma WJ, Bethge M, Tolias AS (2012) A Fast and Simple Population Code for Orientation in Primate V1. J Neurosci 32: 10618–10626.

Bichler O, Querlioz D, Thorpe SJ, Bourgoin JP, Gamrat C (2012) Extraction of temporally correlated features from dynamic vision sensors with spike-timing-dependent plasticity. Neural Networks 32: 339–348.

Birznieks I, Jenmalm P, Goodwin AW, Johansson RS (2001) Encoding of direction of fingertip forces by human tactile afferents. J Neurosci 21: 8222–8237.

Birznieks I, Vickery RM (2017) Spike Timing Matters in Novel Neuronal Code Involved in Vibrotactile Frequency Perception. Curr Biol 27: 1485–1490.e2.

Birznieks I, Wheat HE, Redmond SJ, Salo LM, Lovell NH, Goodwin AW (2010) Encoding of tangential torque in responses of tactile afferent fibres innervating the fingerpad of the monkey. J Physiol 588: 1057–1072.

Bisley JW, Goodwin AW, Wheat HE (2000) Slowly adapting type I afferents from the sides and end of the finger respond to stimuli on the center of the fingerpad. J Neurophysiol 84: 57–64.

Christel MI, Kitzel S, Niemitz C (1998) How Precisely Do Bonobos (Pan paniscus) Grasp Small ObjectsT? Int J Primatol 19: 165–194.

Clay Reid R, Alonso JM, Reid RC, Alonso JM (1995) Specificity of monosynaptic connections from thalamus to visual cortex. Nature 378: 281–284.

Cohen MR, Kohn A (2011) Measuring and interpreting neuronal correlations. Nat Neurosci 14: 811–819.

Delhaye BP, Long KH, Bensmaia SJ (2018) Neural Basis of Touch and Proprioception in Primate Cortex. In: Comprehensive Physiology, pp 1575–1602. Hoboken, NJ, USA: John Wiley & Sons, Inc.

DiCarlo JJ, Johnson KO (1999) Velocity invariance of receptive field structure in somatosensory cortical area 3b of the alert monkey. J Neurosci 19: 401–419.

DiCarlo JJ, Johnson KO (2000) Spatial and temporal structure of receptive fields in primate somatosensory area 3b: effects of stimulus scanning direction and orientation. J Neurosci 20: 495–510.

DiCarlo JJ, Johnson KO (2002) Receptive field structure in cortical area 3b of the alert monkey. Behav Brain Res 135: 167–178.

DiCarlo JJ, Johnson KO, Hsiao SS (1998) Structure of receptive fields in area 3b of primary somatosensory cortex in the alert monkey. J Neurosci 18: 2626–2645.

Dodson MJ, Goodwin AW, Browning AS, Gehring HM (1998) Peripheral neural mechanisms determining the orientation of cylinders grasped by the digits. J Neurosci 18: 521–530.

Friedman RM, Khalsa PS, Greenquist KW, Lamotte RH (2002) Neural coding of the location and direction of a moving object by a spatially distributed population of mechanoreceptors. J Neurosci 22: 9556–9566.

Goodwin AW, Browning a S, Wheat HE (1995) Representation of curved surfaces in responses of mechanoreceptive afferent fibers innervating the monkey’s fingerpad. J Neurosci 15: 798–810.

Goodwin AW, Wheat HE (2002) How is tactile information affected by parameters of the population such as non-uniform fiber sensitivity, innervation geometry and response variability? Behav Brain Res 135: 5–10.

Goodwin AW, Wheat HE (2004) Sensory signals in neural populations underlying tactile perception and manipulation. Annu Rev Neurosci 27: 53–77.

Graczyk EL, Schiefer MA, Saal HP, Delhaye BP, Bensmaia SJ, Tyler DJ (2016) The neural basis of perceived intensity in natural and artificial touch. Sci Transl Med 8:362ra142–362ra142.

Hsiao SS, Lane J, Fitzgerald PJ (2002) Representation of orientation in the somatosensory system. Behav Brain Res 135: 93–103.

Hubel DH, Wiesel TN (1962) Receptive fields, binocular interaction and functional architecture in the cat’s visual cortex. J Physiol 160: 106–154.

Jenmalm P, Birznieks I, Goodwin AW, Johansson RS (2003) Influence of object shape on responses of human tactile afferents under conditions characteristic of manipulation. Eur J Neurosci 18: 164–176.

Johansson RS (1978) Tactile sensibility in the human hand: receptive field characteristics of mechanoreceptive units in the glabrous skin area. J Physiol 281: 101–125.

Johansson RS, Birznieks I (2004) First spikes in ensembles of human tactile afferents code complex spatial fingertip events. Nat Neurosci 7: 170–177.

Johansson RS, Flanagan JR (2009) Coding and use of tactile signals from the fingertips in object manipulation tasks. Nat Rev Neurosci 10: 345–359.

Johansson RS, Vallbo AB (1979) Tactile sensibility in the human hand: relative and absolute densities of four types of mechanoreceptive units in glabrous skin. J Physiol 286: 283–300.

Johnson KO, Hsiao SSS (1992) Neural mechanisms of tactual form and texture perception. Annu Rev Neurosci 15: 227–250.

Khalsa PS, Friedman RM, Srinivasan MA, Lamotte RH (1998) Encoding of shape and orientation of objects indented into the monkey fingerpad by populations of slowly and rapidly adapting mechanoreceptors. J Neurophysiol 79: 3238–3251.

Killebrew JH, Bensmaia SJ, Dammann JF, Denchev P, Hsiao SS, Craig JC, Johnson KO (2007) A dense array stimulator to generate arbitrary spatio-temporal tactile stimuli. J Neurosci Methods 161: 62–74.

LaMotte RH, Lu C, Srinivasan MA (1996) Tactile neutral codes for the shapes and orientation of objects. In: Somesthesis and the Neurobiology of the Somatosensory Cortex (Franzen O, Johansson R, Terenius L, eds), pp 113–122. Basel: Birkhäuser Basel.

LaMotte RH, Srinivasan MA (1987a) Tactile discrimination of shape: responses of slowly adapting mechanoreceptor afferents to a step stroked across the monkey fingerpad. J Neurosci 7: 1655–1671.

LaMotte RH, Srinivasan MA (1987b) Tactile discrimination of shape: responses of rapidly adapting mechanoreceptive afferents to a step stroked across the monkey fingerpad. J Neurosci 7: 1672–1681.

Lechelt EC (1992) Tactile spatial anisotropy with static stimulation. Bull Psychon Soc 30: 140–142.

Loomis M (1980) An Investigation of Tactile Hyperacuityl. Sens Processes 302: 289–302.

Mackevicius EL, Best MD, Saal HP, Bensmaia SJ (2012) Millisecond precision spike timing shapes tactile perception. J Neurosci 32: 15309–15317.

Mountcastle VB, Talbot WH, Darian-Smith I, Kornhuber HH (1967) Neural basis of the sense of flutter-vibration. Science (80-) 155: 597–600.

Muniak MA, Ray S, Hsiao SS, Dammann JFJF, Bensmaia SJ (2007) The neural coding of stimulus intensity: linking the population response of mechanoreceptive afferents with psychophysical behavior. J Neurosci 27: 11687–11699.

Pack CC, Bensmaia SJ (2015) Seeing and Feeling Motion: Canonical Computations in Vision and Touch. PLOS Biol 13:e1002271.

Peters RM, Staibano P, Goldreich D (2015) Tactile orientation perception: an ideal observer analysis of human psychophysical performance in relation to macaque area 3b receptive fields. J Neurophysiol 114: 3076–3096.

Phillips JR, Johnson KO (1981) Tactile spatial resolution. II. Neural representation of Bars, edges, and gratings in monkey primary afferents. J Neurophysiol 46: 1192–1203.

Phillips JR, Johnson KO, Phillips JR (1981) Tactile spatial resolution. III. A continuum mechanics model of skin predicting mechanoreceptor responses to bars, edges, and gratings. J Neurophysiol 46: 1204–1225.

Platkiewicz J, Lipson H, Hayward V (2016) Haptic Edge Detection Through Shear. Sci Rep 6: 1–15.

Pruszynski JA, Flanagan JR, Johansson RS (2018) Fast and accurate edge orientation processing during object manipulation. Elife 7:e31200.

Pruszynski JA, Johansson RS (2014) Edge-orientation processing in first-order tactile neurons. Nat Neurosci 17: 1404–1409.

Pubols LM, Leroy RF (1977) Orientation detectors in the primary somatosensory neocortex of the raccoon. Brain Res 129: 61–74.

Saal HP, Delhaye BP, Rayhaun BC, Bensmaia SJ (2017) Simulating tactile signals from the whole hand with millisecond precision. Proc Natl Acad Sci U S A 114:E5693–E5702.

Saal HP, Wang X, Bensmaia SJ (2016) Importance of spike timing in touch: an analogy with hearing? Curr Opin Neurobiol 40: 142–149.

Srinivasan MA, LaMotte RH (1987) Tactile discrimination of shape: responses of slowly and rapidly adapting mechanoreceptive afferents to a step indented into the monkey fingerpad. J Neurosci 7: 1682–1697.

Sripati AP, Bensmaia SJ, Johnson KO (2006) A Continuum Mechanical Model of Mechanoreceptive Afferent Responses to Indented Spatial Patterns. J Neurophysiol 95: 3852–3864.

Suresh AK, Saal HP, Bensmaia SJ (2016) Edge orientation signals in tactile afferents of macaques. J Neurophysiol 116: 2647–2655.

Talbot WH, Darian-Smith I, Kornhuber HH, Mountcastle VB (1968) The sense of flutter-vibration: comparison of the human capacity with response patterns of mechanoreceptive afferents from the monkey hand. J Neurophysiol 31: 301–334.

Thorpe SJ, Delorme A, Van Rullen R (2001) Spike-based strategies for rapid processing. Neural Networks 14: 715–725.

Van Rullen R, Guyonneau R, Thorpe SJ (2005) Spike times make sense. Trends Neurosci 28: 1–4.

Victor JD, Purpura KP (1997) Metric-space analysis of spike trains: theory, algorithms and application. Netw Comput Neural Syst 8: 127–164.

Weber AI, Saal HP, Lieber JD, Cheng J-W, Manfredi LR, Dammann JF, Bensmaia SJ (2013) Spatial and temporal codes mediate the tactile perception of natural textures. Proc Natl Acad Sci U S A 110: 17107–17112.

Wheat HE, Goodwin AW, Browning AS (1995) Tactile resolution: peripheral neural mechanisms underlying the human capacity to determine positions of objects contacting the fingerpad. J Neurosci 15: 5582–5595.

Yau JM, Pasupathy A, Fitzgerald PJ, Hsiao SS, Connor CE (2009) Analogous intermediate shape coding in vision and touch. Proc Natl Acad Sci U S A 106: 16457–16462.

